# Mesenchymal stem cells of the bone marrow raise infectivity of *Plasmodium falciparum* gametocytes

**DOI:** 10.1101/2023.08.08.552490

**Authors:** Ragavan Varadharajan Suresh, Bingbing Deng, Yonas Gebremicale, Kyle Roche, Kazutoyo Miura, Carole Long

**Affiliations:** Laboratory of Malaria and Vector Research, National Institute of Allergy and Infectious Diseases, National Institutes of Health, Rockville, Maryland, USA

**Author notes:** Address correspondence(s) to Carole Long, or Kazutoyo Miura.

**Keywords:** Malaria, *Plasmodium falciparum*, gametocytes, mesenchymal stem cells, bone marrow, mosquito infectivity

## Abstract

*Plasmodium falciparum* is a parasite that causes the deadly human disease, malaria, and exhibits a complex life cycle in the human and mosquito hosts. As the sexual stages of the parasite, gametocytes mature in the human body and propagate malaria when they are picked up by mosquitoes to infect new hosts. Previous research has shown that gametocytes home to the bone marrow of the host where they complete their maturation and alter the behavior of resident Mesenchymal Stem Cells (MSCs). In this study, we investigated the alternate side of this host-pathogen interaction, whether MSCs could alter the behavior of gametocytes. Gametocytes were co-cultured with MSCs until maturity and subsequently fed to mosquitoes to measure the oocysts produced. Here we report for the first time, that MSCs co-culture significantly elevated oocyst numbers in the infected mosquito compared to conventional culture medium. This enhancement appeared to be most effective during the early stages of gametocyte development and was not replicated by other cell types. MSC co-culture also increased infectivity of field isolated *P. falciparum* parasites. This effect was partially mediated by soluble factor(s) as conditioned medium harvested from MSCs could also partially raise infectivity of gametocytes to nearly half compared to MSC co-culture. Together this study reveals novel host pathogen interactions, where the human MSCs are elevating the infectivity of malaria gametocytes.

**Importance:** While prior research has established that *Plasmodium* gametocytes sequester in the bone marrow and can influence resident stem cells, the question of why they would choose this compartment and these cells remained a mystery. This study for the first time, shows that being in the presence of MSCs alters the biology of the *P. falciparum* parasite and makes it more infectious to mosquitoes, hinting at novel mechanisms in its life cycle. This method also facilitates mosquito infections with field isolated parasites, affording research teams new infection models with parasite s which are challenging to infect into mosquitos using conventional culture methods. Finally, our findings that MSC conditioned medium can also raise infectivity opens avenues of investigation into mechanisms involved, but can also serve as a practical tool to researchers hoping to increase oocyst yields.

## Introduction

Malaria is one of the oldest and deadliest diseases plaguing humanity with tremendous infectivity caused by the protozoan parasites of the genus *Plasmodium* with five species infecting humans *P. falciparum, P. vivax, P. ovale, P. malariae and P. knowlesi* (1, 2). The focus of this study is the parasite *P. falciparum* which infects a human host through the bite of an infected mosquito and has a brief developmental period in the liver before beginning its erythrocytic life cycle (3, 4). A large percentage of the parasites in the erythrocytes replicate asexually, increasing parasite numbers, causing RBC lysis, clogging small blood vessels and contributing to most of the pathology associated with malaria (3, 5, 6). Over time, a small percentage of the erythrocytic parasites deviate from the asexual replicative cycle to follow the path of sexual reproduction and are referred to as gametocytes (7). These mature male and female gametocytes are picked up by mosquitos to perpetuate infection.

Gametocytes of *P. falciparum* require around eleven days to mature and go through five morphologically distinct stages (Stage I – V) (8–10). Reports dating as far back as 1900s (11) observed only mature stage V gametocytes in the peripheral circulation while the earlier stages were sequestered in the spleen or bone marrow (12, 13). Interestingly, the bone marrow has the highest ratio of gametocytes over asexual parasites compared to those in other organs, whether the infection is with *P. falciparum, P. vivax,* or *P. berghei* (14–18). But why this particular affinity for the bone marrow and what resident cells could be key players? Unlike asexual stages, gametocytes do not bind endothelial cells and are primarily found in the extravascular compartment of the bone marrow (15, 19). Here, they are enriched around erythroblastic islands, developing inside erythroblasts and reticulocytes (14, 20, 21). One explanation is the rigidity of early stage gametocytes that allows them to be retained in the fine vasculature of the marrow and become more deformable as they mature and egress from the marrow (22, 23). In addition, a recent study by Messina et al showed that immature gametocyte infected red blood cells bind to bone marrow mesenchymal stem cells (MSCs) (24).

MSCs are pluripotent stem cells in the bone marrow and can differentiate into osteoblasts, myocytes, adipocytes, or chondrocytes depending on the needs of the body. They are minimally defined as cells which are (a) plastic-adherent, (b) express CD105, CD73, and CD90 on their surface, while lacking expression of CD45, CD34, CD14, CD79alpha, HLA-DR and (c) able to differentiate into osteoblasts, adipocytes, and chondrocytes in vitro (25). They aid in routine cellular regeneration as well as healing and replenishing damaged tissues during disease states (26, 27), the full extent of which is still under active investigation. While the specific ligand(s) and receptor(s) for the binding of immature gametocytes and MSCs were not identified in the Messina study, the bound gametocytes were reported to induce the secretion of pro-angiogenetic factors from the stem cells. However, the study did not investigate if the host cells were altering the behavior of parasites. Around the same time, a report by Chen et al showed that MSCs could prolong the lifespan of RBCs in storage, as a type of cellular preservative (28). Given the complexity of the parasite and its success as a pathogen, it seemed likely that this sequestration in the bone marrow and interaction with MSCs might have an impact on the nature of the gametocyte. We hypothesized that the presence of MSCs might raise the fitness of *P. falciparum* gametocytes. While this increase in fitness may manifest in various ways, we evaluated the mosquito infectivity (number of oocysts) in mosquitoes, which were fed gametocytes co-cultured with or without MSCs, as a primary measure.

Our results show that these co-cultured gametocytes yielded markedly higher oocyst numbers in the midguts of the infected mosquitos relative to medium controls. We also show that medium conditioned by MSCs had an enhancement effect on gametocytes. Our work is the first to prove that MSCs elevate mosquito infectivity of *P. falciparum* gametocytes.

## Results

### Co-culturing *P. falciparum* NF54 gametocytes with mesenchymal stem cells (MSCs) raised infectivity to mosquitos

To investigate biological significance of gametocytes being in the presence of human MSCs, *P. falciparum* gametocytes were co-cultured (CC) in the presence of MSCs for eleven days to generate mature gametocytes. Given the existing reports detailing the importance of TNFα induced ligands on the cytoadherence of asexual stage parasites to endothelial cells (29), MSCs were pre-treated with 1ng/mL of TNFα overnight prior to starting co-culture (MSC CC+TNFα), to allow for the expression of any ligands which could potentially affect gametocyte development. Gametocytes were also cultured in two medium controls (conventional gametocyte culture medium (Conv), and hybrid culture medium, (Hyb)). The hybrid culture medium was a 1:1 mixture of conventional gametocyte culture medium and MSC culture medium. This Hyb was used in all co-culture conditions described in this study instead of 100% conventional gametocyte culture medium or MSCs culture medium. Stage V gametocytemia and exflagellation were measured for all cultures on the day of feed. This was to evaluate the effect of MSC CC on gametocyte development and activity relative to established Conv conditions. The stage V gametocytemia (%)/exflagellation (per µL of 25% hematocrit concentrated culture) for Conv, Hyb, and MSC CC+TNFα were 1.94/1285, 2.98/620, and 1.82/1000, respectively (Fig. 1A). This indicated that gametocyte development was similar in Conv and MSC CC+TNFα. Representative images of fixed, Giemsa-stained gametocytes from gametocytes cultured in Conv (Fig. 1E), Hyb (Fig. 1F) and MSC CC+TNFα (Fig. 1G) are shown, and gametocytes in all three conditions appeared morphologically similar. The gametocytes from the 3 conditions were diluted to 0.4% stage V gametocytemia and fed to mosquitoes. Eight days after this feed, the mosquitos were dissected and oocysts in their midgut were enumerated. If the presence of MSC cells was beneficial to the gametocytes, then we expected to see an increase in oocyst numbers relative to the medium controls and vice versa. MSC CC+TNFα (average of 172 oocysts per midgut) had significantly higher oocysts (*P =* 0.002) compared to Conv control, and there was no significant difference between Conv and Hyb (41 in Conv and 34 in Hyb; Fig. 1B). To confirm our initial observation, an independent assay was conducted with the same three conditions, but the final gametocytemia was adjusted to 0.1% for the feeding experiment. While Stage V gametocytemia and exflagellation were not obviously higher in MSC CC+TNFα compared to Conv (Fig. 1C), again MSC CC+TNFα (average of 77) led to significantly higher (*P =* 0.002) oocysts per midgut compared to Conv (average of 9, Fig. 1D). There was no significant difference in oocyst number between Conv and Hyb in the second experiment as well. The two experiments confirmed that MSC CC+TNFα could induce higher oocysts compared to Conv without increasing Stage V gametocytemia or exflagellation. Since there was no difference in oocyst numbers between Conv and Hyb in the two experiments, the following studies were conducting using Hyb as medium control, which used the same medium as MSC CC+TNFα conditions.

**FIG 1:**
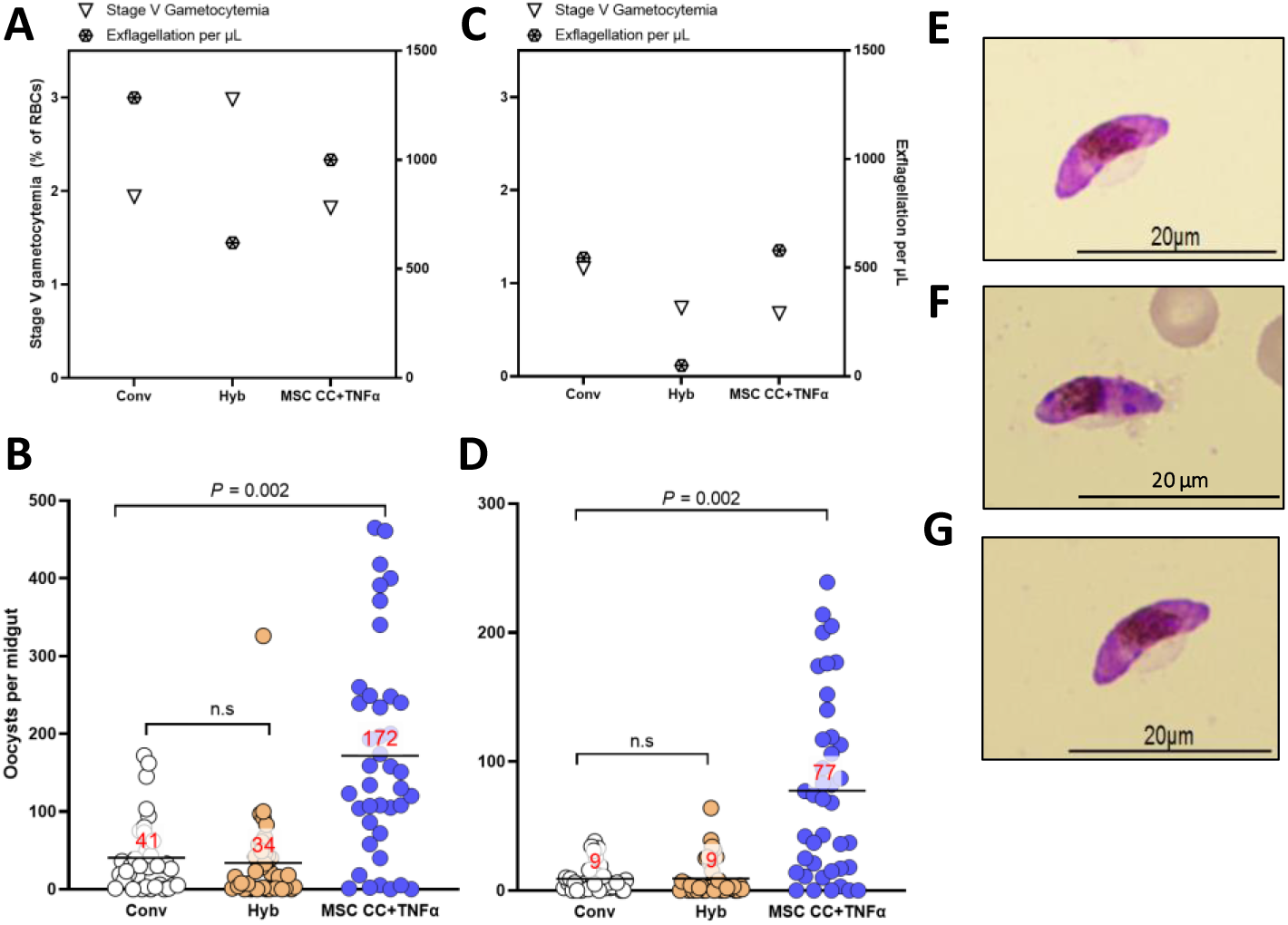
MSC Co-culture raised infectivity of malaria gametocytes. *P. falciparum* NF54 gametocytes were co-cultured with human mesenchymal stem cells, which were pre-treated overnight with 1ng/mL TNFα, (MSC CC+TNFα) for eleven days and used to infect mosquitos. Gametocytes raised in conventional gametocyte culture medium (Conv) and hybrid culture medium (Hyb) served as medium controls. Corresponding percent Stage V gametocytemia and exflagellation per microliter post co-culture are shown (A). These groups were fed to mosquitos and resulting oocysts per midgut in individual mosquitoes (circles) are shown with the arithmetic mean (bars with red text) (B). An independent experiment was conducted for the same three conditions, and the results are shown in (C) and (D). *P* denotes p-values calculated by a zero inflated negative binomial (ZINB) model with a Bonferroni correction. n.s.; not significant. Representative light microscopy images of fixed, Giemsa-stained gametocytes post culture are presented for Conv (E), Hyb (F) and MSC CC+TNFα (G).

### Pre-treating MSCs with TNF alpha was not required for conferring elevating gametocyte infectivity

To investigate whether the TNFα pre-treatment used in Fig 1 was necessary, MSC co-culture was performed with or without pre-treatment. After infecting mosquitoes, gametocytes co-cultured with MSCs in the absence of TNFα (MSC CC) yielded an average of 67 oocysts per midgut and gametocytes co-cultured with TNFα pre-treated MSCs (MSC CC+ TNFα) yielded an average of 63 oocysts per midgut, a statistically insignificant difference (*P =* 0.84). Importantly, the MSC CC group was still significantly higher than the Hyb control’s 3 oocysts per midgut (*P =* 0.002; Fig 2A). In addition to this experiment, a total of 10 independent experiments were performed without the TNFα pre-treatment and MSC CC always showed higher oocysts than Hyb in each experiment (p=0.002;Fig 2B). These results confirmed that the TNFα pre-treatment was not required to induce higher infectivity by MSC CC. Thus, the subsequent experiments were conducted without TNFα pre-treatment.

**FIG 2:**
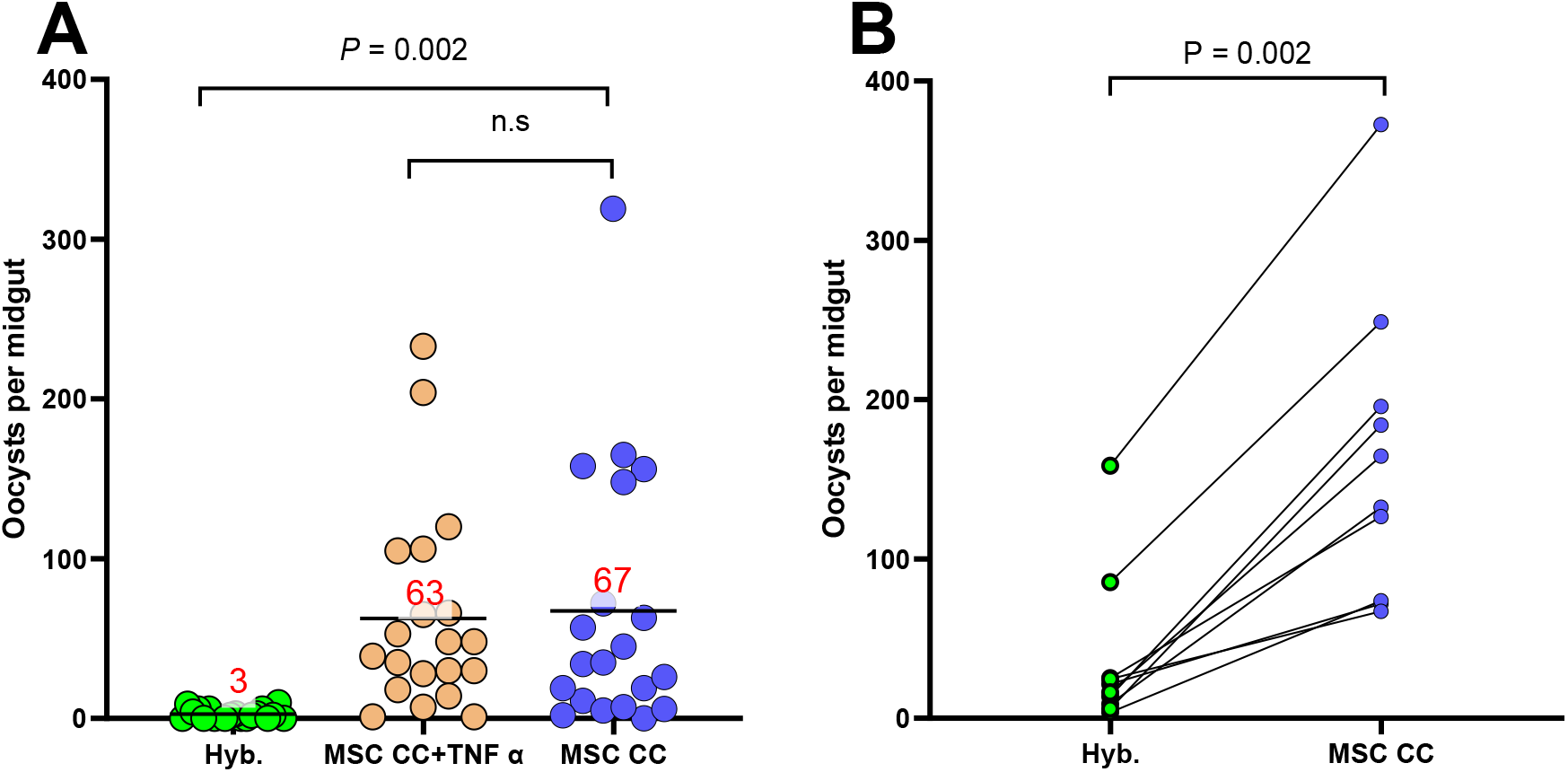
Pre-treatment of MSCs with TNFα was not required to confer increased gametocyte infectivity. MSCs treated with 1ng/mL of TNFα overnight (MSC CC+TNFα) or non-treated MSCs (MSC CC) were used to co-culture *P. falciparum* NF54 gametocytes for eleven days. Oocysts per midgut are shown with the arithmetic means (bars with red text)(A). Data from ten additional independent experiments comparing average oocysts from gametocytes cultured in control media (Hyb) versus co-cultured with MSCs that didn’t receive pre-treatment with TNFα (MSC CC) are shown. The connected lines indicate oocyst data collected from the same experiment (B). *P* values in (A) denote p-values calculated by ZINB model with a Bonferroni correction. n.s.; not significant. p value in (B) was calculated by a Wilcoxon matched pairs signed rank test.

### A similar increase in infectivity was not conferred by three other tested cell types or coating culture flasks with substrates

The next question was whether the increase of infectivity was unique to MSCs, or if other cell types or organic basement substrates (which can support cell growth in vitro (30, 31)) could also raise infectivity of gametocytes. The first was investigated by co-culturing gametocytes with Human Umbilical Vein Endothelial Cells (HUVEC), human GH354 cells (a cervical epithelial cell line), and murine Distal Convoluted Tubule cells (mDCT). Like MSCs, these are all adherent cells and form a confluent monolayer. Co-culture with HUVEC cells led to 35 oocysts per midgut (Fig 3A), which was slightly higher than MSC Hyb control (21 oocysts per midgut), but less than half of the oocysts in the MSC CC group (72 oocysts per midgut). While the difference between HUVEC CC and MSC CC did not reach statistical significance (P=0.17), the HUVEC Hyb condition yielded zero oocysts, indicating that the medium was toxic to gametocytes, unlike the case with MSC Hyb (Fig 1). When infecting mosquitos, the GH354 co-cultured gametocytes yielded 1 oocyst per midgut, significantly lower than MSC CC’s 373 (p *=* 0.001, Fig 3B). Similarly, mDCT co-cultured gametocytes resulted in 0 oocysts per midgut relative to MSC CC’s significantly higher 72 (*P =* 0.002, Fig 3C). Thus, co-cultures with other cell types were unable to produce the same levels of elevated oocyst numbers produced by MSC CC.

**FIG 3:**
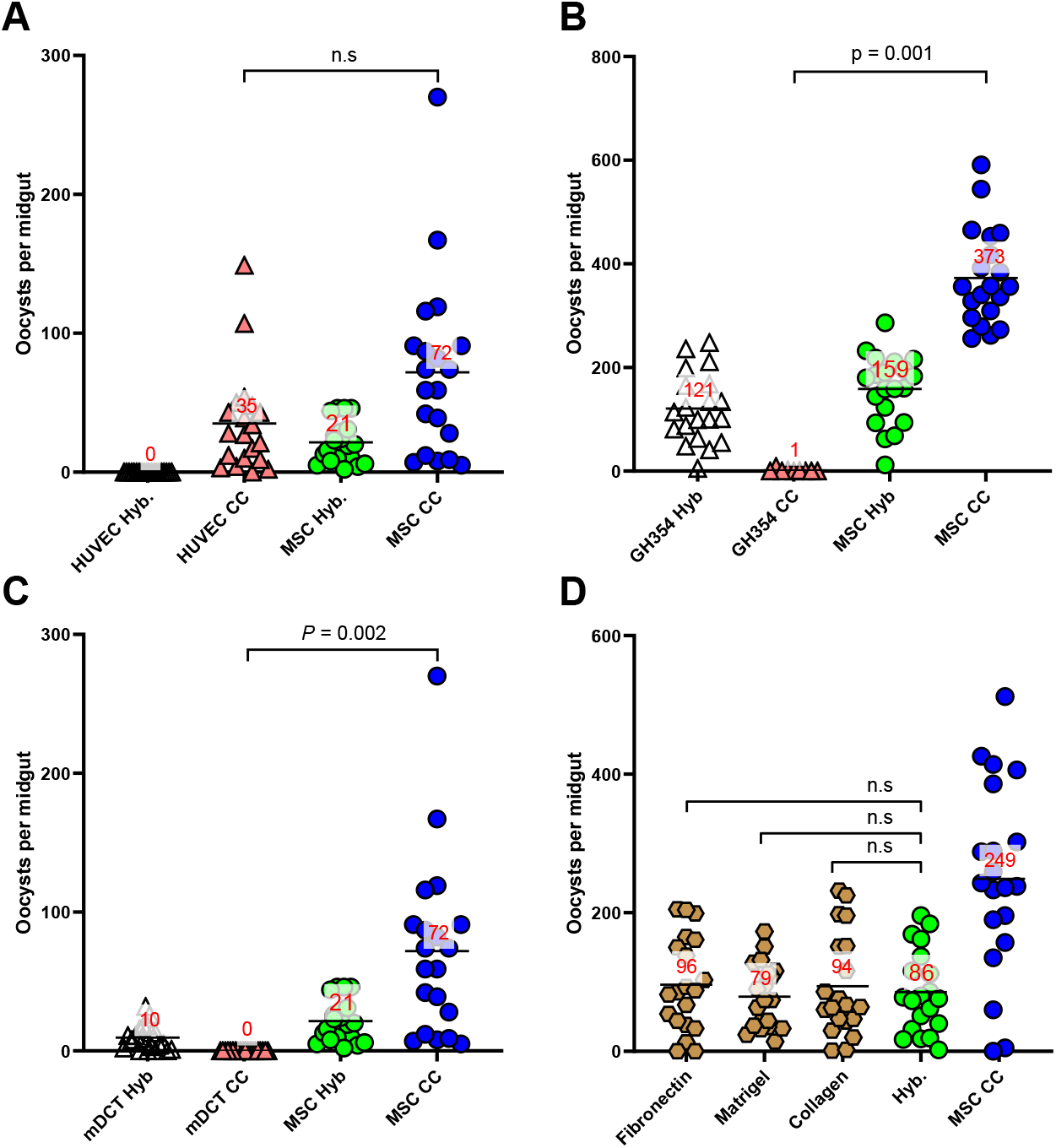
A similar increase in infectivity was not conferred by three other tested cell types or coating culture flasks with substrates. *P. falciparum* NF54 gametocytes were co-cultured with HUVEC (A), GH354 (B) or mDCT cells (C) with corresponding Hyb controls, negative (MSC Hyb) and positive (MSC CC) controls. After 11 days of co-culture, gametocytes were used to infect mosquitos and resulting oocysts per midgut are presented. Independently, gametocytes were cultured in flasks coated with fibronectin, Matrigel, or collagen, for eleven days, used to infect mosquitos and resulting oocysts are presented (D). Bars with red text denote arithmetic means and *P* denote p-values calculated by ZINB model with a Bonferroni correction. n.s.; not significant.

Next, culture flasks were coated with 2.5 μg/cm^2^ fibronectin, 1:100 Matrigel, or 9 μg/cm^2^ collagen. Gametocytes cultured in these coated flasks led to 96, 79, and 94 oocysts, respectively which were similar to Hyb’s 86 (all unadjusted P values of substrate vs Hyb >0.75; all adjusted P values = 1). MSC CC yielded 249 oocysts, over twice as much as any of the other conditions (Unadjusted P values of MSC CC vs substrates range between 0.013 and 0.038; adjusted P values between 0.052 and 0.15, Fig 3D). Thus, coating the plasticware with these substrates did not raise infectivity beyond levels of Hyb control.

### The first seven days of co-culture with MSCs elevated infectivity on par with full duration co-culture

The effect of co-culture duration on oocyst yield was investigated by stopping co-culture after D6 or D10, with full duration (D3 to D14) Hyb and MSC CC serving as controls. After the stipulated co-culture duration, gametocytes were transferred to flasks without MSCs, then raised in Hyb medium until D14. When these cultures were used to infect mosquitoes, Full Hyb, D3-6, D3-10, and Full MSC CC led to 55, 168, 300 and 281 oocysts, respectively (Fig 4). We observed that as little as four days of co-culture (D3-6) was able to raise infectivity beyond Full Hyb control (*P =* 0.02). Maintaining co-culture for an additional four days yielded 300 oocysts, nearly double that from D3-6. While this four-day increase failed to reach statistical significance when compared to D3-6 (*P =* 0.28), it was on par with full duration co-culture (*P =* 1). Thus, the first seven days of co-culture were sufficient to confer infectivity on par with full duration co-culture, and the first four days were sufficient to confer more than medium control. The effect of co-culture on the final four days of gametocyte development (D11-14) was also tested and this condition yielded 97 oocysts, which was in the middle of Hyb (55) and D3-6 (168) (Fig S1).

**FIG 4:**
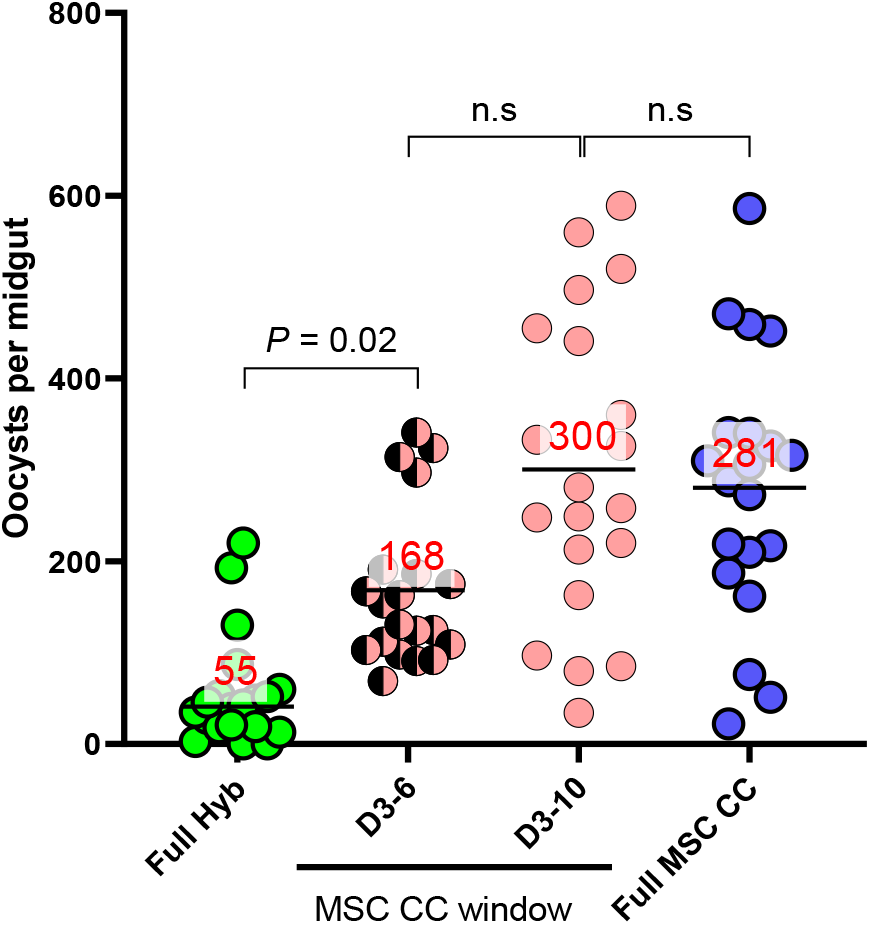
The first seven days of co-culture with MSCs elevated infectivity on par with full duration co-culture. *P. falciparum* NF54 gametocytes were co-cultured with MSCs for D3-6, D3-10, and full duration (D3-14, Full MSC CC) or full duration Hyb. These gametocytes were then used to infect mosquitos and resulting oocysts per midgut are presented with the arithmetic means (bars with red text). *P* denote p-values calculated by ZINB model with a Bonferroni correction. n.s.; not significant

### Co-culture with MSC raised infectivity of field isolated parasites

To assess whether this novel observation was limited to the NF54 strain, which has been maintained in laboratory culture for decades, two parasite lines isolated from Cambodia (Cambodia #887 and #806) and other two lines from Mali (KN #1068 and #1254) were co-cultured with MSCs. To maximize the chance of observing elevated oocyst yields, the study was carried out for full-duration co-culture. In keeping with prior observations, MSC co-culture of field parasites led to significantly elevated oocyst numbers compared to Hyb controls in three out of four isolates tested (p=0.001 for Cambodia # 887, KN # 1068 and KN # 1254 in Fig 5A, 5C and 5D, respectively); the increase in the other isolate (Cambodia # 806) followed the same trend but failed to reach statistical significance (P=0.058, Fig 5B). This revealed that the observed phenomena were not limited to the laboratory adapted NF54.

**FIG 5:**
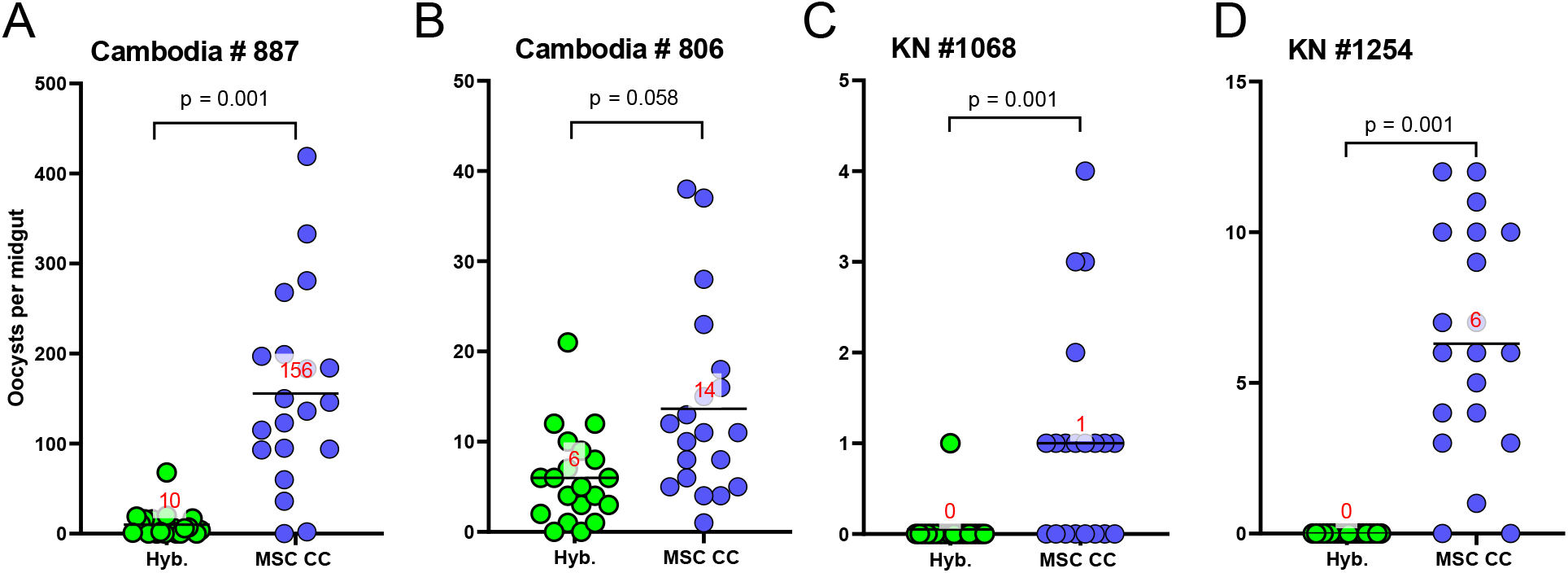
Co-culture with MSC raised infectivity of field isolated parasites. Two field isolates each of *P. falciparum* from Cambodia (A,B) and Mali (C,D) were co-cultured with MSC cells alongside Hyb media controls. Oocysts per midgut are shown with the arithmetic means (bars with red text) and p-values were calculated by ZINB model.

### Medium conditioned by stem cells raised infectivity of gametocytes

Since being in the presence of MSCs was elevating infectivity, we sought to determine if contact was required or the environment around the MSCs was elevating infectivity.

MSCs were grown to confluence and incubated for 24 hours in Hyb medium in flasks referred to as “feeder flasks”. After incubation, the culture supernatant from feeder flasks was collected, filtered, and used as a conditioned medium (CM). When gametocytes cultured in CM were used to infect mosquitos, they led to significantly higher oocysts than the Hyb medium control (32 oocysts from CM versus 4 in Hyb, *P =* 0.004), albeit less than half of MSC CC (74 oocysts *P =* 0.09; Fig 6A). To investigate if the CM could be made better, in the next experiment three different feeder flasks were setup. A flask containing MSCs alone (MSC^+^iRBC^-,^ same as CM above), another containing infected RBCs alone (MSC^-^iRBC^+^ CM), and finally a co-culture of MSCs and infected RBCs (MSC^+^iRBC^+^ CM). The iRBCs contained a mix of synchronized gametocytes as well as asexual stage parasites (the same iRBCs used for the MSC CC). CM was harvested and filtered every 24 hours from each of these feeders and used to raise gametocytes (alongside MSC CC and Hyb control gametocytes). After 11 days of culture, gametocytes raised in these conditions were used to infect mosquitos. We observed that gametocytes raised in CM harvested from the co-culture (MSC^+^iRBC^+^ CM; 16 oocysts) and from iRBCs alone (MSC^-^iRBC^+^ CM; 10 oocysts) led to significantly lower oocysts than Hyb condition (65 oocysts, P=0.002 in both comparisons; Fig 6B). Next, to rule out effect of harmful products which could be generated by the infected RBCs, CM was generated from feeder flasks containing MSCs co-cultured with uninfected RBCs (uRBC). Culturing gametocytes in this CM from MSC^+^uRBC^+^ did not alter their infectivity relative to CM from MSCs alone (29 oocysts for MSC^+^uRBC^+^ CM versus 33 in MSC^+^RBC^-^ CM, *P =* 0.76; Fig 6C). This revealed that MSCs conditioned the medium independent of whether iRBCs (or uRBCs) were in physical contact. To support future identification of the responsible molecule(s) in the conditioned medium (CM from MSCs alone), we compared fresh (unfrozen) CM and CM subjected to one-time freeze-and-thaw. The frozen and thawed CM yielded 50 oocysts, higher than Hyb control’s 9 (*P* = 0.002) and was on par with fresh CM’s 63 (*P =* 1.0; Fig 7A). Again, the frozen and thawed CM resulted in about half of the oocysts seen with the MSC CC. An independent experiment showed similar results (Fig 7B). Thus, CM can be stored in a frozen state and retain potency after thawing for at least one round.

**FIG 6.**
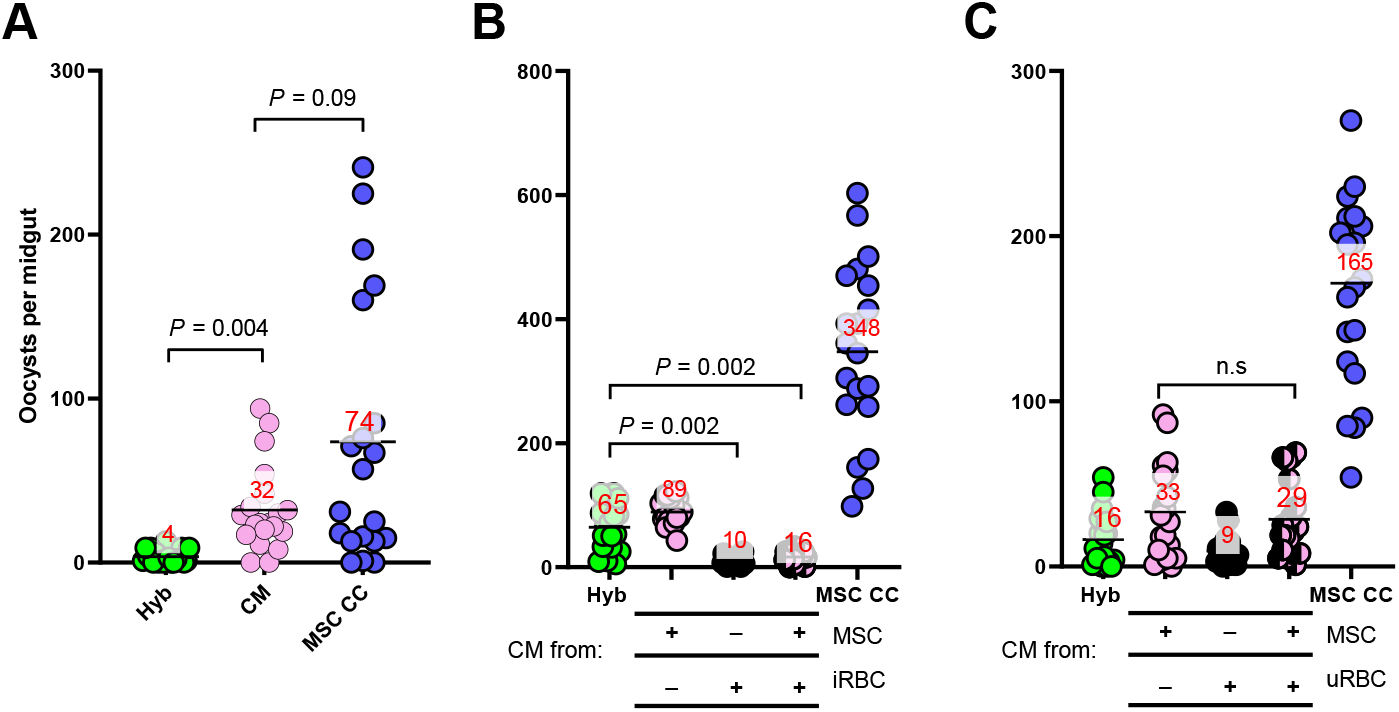
Medium conditioned by stem cells raised infectivity of gametocytes. MSCs were grown to confluence and incubated for 24 hours in Hyb medium and after incubation, the culture supernatant was collected, filtered, and used as a conditioned medium (CM). *P. falciparum* NF54 gametocytes were cultured in the CM, or in the control conditions (Hyb and MSC CC) (A). Independently, CM were prepared from 3 different conditions; MSC^+^iRBC^--^(the same condition as panel A), MSC^-^iRBC^+^, or MSC^+^iRBC^+^, where iRBC denotes infected red blood cells (RBCs). Gametocytes were cultured with one of the three CM alongside Hyb and MSC CC controls (B). In the next experiments, CM were prepared in the presence or absence of uninfected RBCs (uRBC), then gametocytes were cultured as B (C). After 11 days of culture, all gametocytes were fed to mosquitos and resulting oocysts per mosquito are presented with the arithmetic mean (bars with red text). *P* denote p-values calculated by ZINB model with a Bonferroni correction. n.s.; not significant

**FIG 7.**
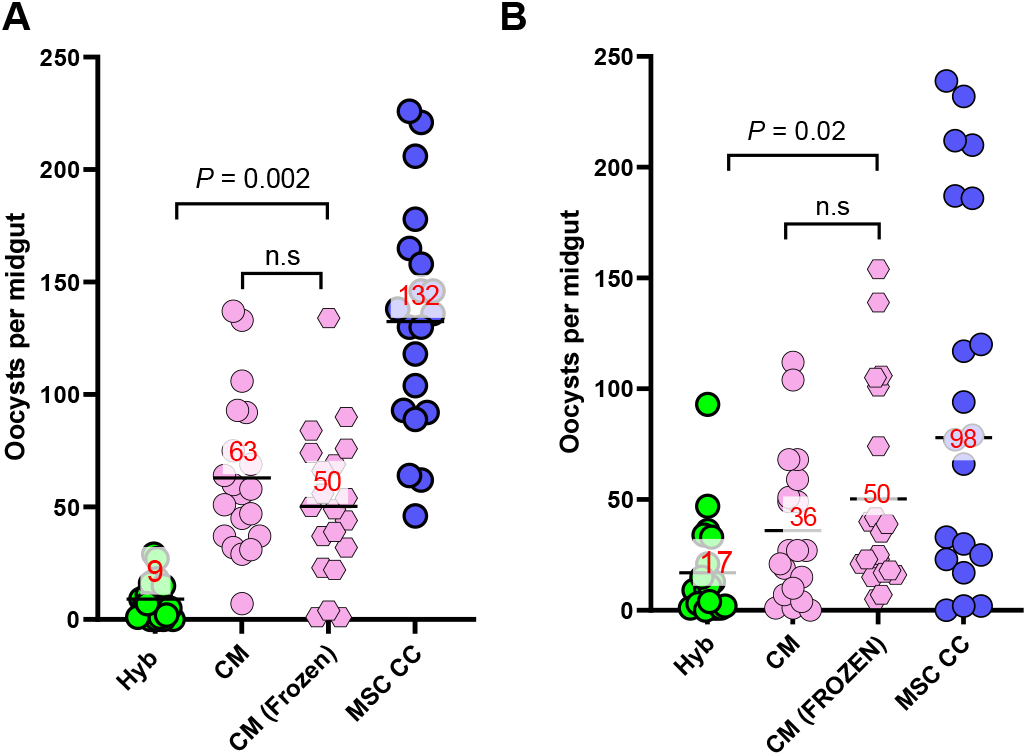
Conditioned medium can be stored frozen. *P. falciparum* gametocytes were cultured in freshly harvested CM (MSCs alone), or CM that was previously frozen and thawed one time, along with Hyb and MSC CC controls (A). After 11 days of culture, these gametocytes were used to infect mosquitos and resulting oocysts per midgut are presented with the arithmetic means (bars with red text). An independent experiment was performed with the same conditions (B) *P* denotes p-values calculated by ZINB model with Bonferroni corrections. n.s.; not significant.

## Discussion

For nearly 130 years malaria gametocytes have been observed in the bone marrow (32), a complex environment of erythropoietic and mesenchymal precursor cells producing numerous other cell types (33, 34). Understandably, significant attention has been paid to the interactions of malaria, a blood borne pathogen, with the erythropoietic compartment of marrow (20, 21). Existing research has shown that gametocytes in the bone marrow delay erythropoietic development, contribute to anemia in the infected individuals (20, 35), and stimulate secretion of pro-angiogenetic factors from the mesenchymal stem population (24). Our study is the first to reveal that host cells, specifically MSCs, can affect the biology of *P. falciparum* gametocytes in ways that are beneficial to transmission.

This discovery provides a new perspective on the previously understood antagonistic relationship between host cells and gametocytes, in which host cells, particularly the immune system, sought to eliminate gametocytes. Now, it is clear that there is a more complex dynamic at play, and gametocytes become more infectious when in the presence of MSCs. This is not limited to NF54, the increased infectivity by MSC co-culture was also observed in four parasite lines isolated from patients in Cambodia and Mali (with Cambodia #806 falling just short of statistical significance). Particularly striking were the observations that post MSC co-culture, the Malian parasites, even with undetectably low gametocytemia and exflagellation, were able to infect mosquitos (Table S1). Perhaps malaria sequestering in the patient bone marrow may be obtaining the same kind of increase in infectivity that is observed in this ex vivo study, partially explaining why patients with submicroscopic gametocytemia can still infect mosquitoes in the field (36, 37).

*P. falciparum* NF54 has been instrumental in understanding malaria infections since it reliably makes large numbers of oocysts in mosquito infections unlike other strains (38, 39). Furthermore, almost all gametocyte biological knowledge for *P. falciparum* malaria has originated from studies using NF54, the clone 3D7, or transgenic parasites made from NF54/3D7. Not only would replicating this co-culture procedure with other lines of parasites serve as an important confirmation that the phenomenon is not specific to the NF54 strain but could also pave the way for other research teams to work with and infect field isolated parasites, a typically arduous task more easily.

To determine whether the increase of infectivity by the co-culture was a specific phenomenon due to the MSC, or the results of a general modification of culture conditions, two types of experiments were conducted: co-culture with other cell types and coating culture flasks with various adhesion promoting substrates that can support the growth of other cell types (30, 40, 41). None of the other cells tested, HUVEC (human), GH354 (human) or mDCT (murine) utilized, could produce a similar elevation in infectivity observed in the MSC co-culture. While co-culture with HUVECs did elevate infectivity over the HUVEC hybrid medium, it was clear that the medium was being toxic to the parasites. There are potentially avenues of HUVEC co-culture worth investigating, however, we did not in this study because it is challenging to identify new formulations of medium that could support growth of HUVEC while also being non-toxic to parasites. In addition to co-culture with other cells, gametocytes cultured in substrate coated flasks fared no better than Hyb controls in terms of the ensuing oocysts. While this is by no means an exhaustive screening of all potential cell types and culture conditions available, given the marked elevation observed with MSCs, we chose to focus our efforts on this cell type.

To uncover the underlying mechanism of increased infectivity, our first approach was to utilize Transwell® chambers with gametocytes in one chamber and MSCs in the other, bathed in a common Hyb medium. The experiment would evaluate the need for contact of gametocyte with the MSCs. However, for unknown reasons, these chambers did not support the growth of gametocytes even in the Hyb condition (data not shown), thus a different method of generating conditioned medium was adopted, using conventional culture flasks followed by filtration. As shown in Figs 6 and 7, the CM induced significantly higher oocyst numbers than Hyb control. However, the culture with CM only gave ∼ half of oocysts compared to MSC CC in all three independent experiments. These results indicate that CM, at least the way collected in the current method, cannot completely replicate MSC CC condition. The presence of uninfected RBCs did not alter CM’s elevation of infectivity, and the presence of iRBCs led to the generation of worse conditioned media. This is potentially due to the buildup of parasite waste products or consumption of beneficial molecules produced by the MSCs. Further investigation is required to fully uncover the mechanism by which MSC CC elevates infectivity. Although CM may not be able to explain all, the results that fresh CM and frozen CM gave similar numbers of oocysts (Fig 7) opens the door for long term storage and analysis of a consistent batch of CM.

The basal MSC medium is supplemented with human Insulin like Growth Factor 1 (IGF-1) and human Fibroblast Growth Factor (FGF-b), which could be benefiting gametocyte infectivity. However, there was no significant difference in oocyst numbers between gametocytes raised in Hyb and Conv Medium. Furthermore, in all experiments, the MSC CC condition consistently yielded higher oocysts than Hyb condition, even though both were maintained using the same medium. Thus, such growth factors in the medium are not likely to explain the elevated infectivity. However, MSCs do secrete growth factors such as Vascular Endothelial Growth Factor (VEGF), Hepatocyte Growth Factor (HGF), Fibroblast Growth Factor 2 (FGF2), Angiopoetin-1 (Angpt-1)(27). Further investigation is required whether any such growth factors contribute to the elevated infectivity in MSC CC and/or CM.

Prior research has revealed positive associations between exflagellation values and oocyst density (42). However, in our study, the co-culture with MSC (MSC CC) induced significantly higher oocysts without a dramatic increase in either stage V gametocytemia or exflagellation numbers when compared to Conv condition (Fig. 1). Furthermore, no differences in gross morphology of the parasite were observed via Giemsa staining. Independently, we also showed that gametocytes in MSC CC went through similar developmental stages as compared to the medium controls (Fig S2). This implies that elevated oocyst numbers by MSC co-culture or conditioned medium were due to changes in the underlying biology of the parasite. MSC co-culture during the early stages of gametocyte development (D3-6) had a greater impact on oocyst numbers than later stages of development (D11-14), and D3-10 co-culture was comparable to full (D3-14) co-culture. We suspect the lack of statistical significance when comparing D3-10 vs D3-6 co-culture is more due to the large variability in oocyst numbers among mosquitoes within a group than a lack of phenotype. Further studies are needed to understand the changes occurring in early-stage gametocytes in the presence of MSCs. Expanding on these promising initial findings, transcriptomic analyses of the gametocytes in co-culture and proteomic analysis of the CM will be employed to further unravel the mechanistic drivers at play.

This study has several limitations. Because each experiment requires nearly one month of time (∼3 weeks of ex vivo cultures and ∼1 week post feeding mosquitos), in most cases, only a single experiment was conducted per test condition. Therefore, it is difficult to tell whether small, statistically insignificant differences between conditions indicate that there is no difference between the groups or there is a small but true difference that could be confirmed by repeated independent testing. Second, due to the general difficulty of gametocyte cultures with field parasites, the effect of conditioned medium on field parasites is yet to be evaluated. Third, at this moment we lack sufficient evidence to speculate mechanism(s) of the elevated infectivity by CM. MSCs might secrete “beneficial” molecule(s) to the CM, consume/detoxify “harmful” molecule(s) in the Hyb medium, or combinations of both. Depending on the underlying mechanism(s), changing the assay conditions (e.g., increasing the frequency of medium changes) may or may result in better or worse infectivity. In addition, this study only utilized immortalized MSCs, not primary MSCs due to the challenge in procuring such cells. Further investigations are required to determine if primary MSCs also increase the infectivity of gametocytes. Finally, despite the myriad of cells existing in the bone marrow, we have only evaluated the effect of MSC on the gametocytes. It is exciting to ponder whether co-culture with other cells from the bone marrow could alter the infectivity of gametocytes or whether such phenomenon could be observed in a whole bone marrow 3-D model.

Since many species of *Plasmodium* display preferential sequestration in the bone marrow (14–18), the increased gametocyte infectivity by MSC observed in this study could be occurring in the human body as well and reveal unknown aspects of the malaria life cycle. An immediate application for the novel findings is to utilize MSC co-culture or conditioned medium to generate large amounts of oocysts, infect more mosquitoes with a smaller amount of culture, or possibly generate more sporozoites than obtained by conventional culture methods. The applicability with field parasites is also immediately apparent. Once the underlying mechanism of the increased infectivity by the co-culture or conditioned medium is uncovered, it will allow a better understanding of gametocyte development, create new avenues of basic research, and hopefully design better drugs to combat the transmission of malaria.

## Materials and methods

### Mammalian cell culture

Immortalized human mesenchymal stem cells were purchased from ATCC (Cat: PCS-500-012) and grown per manufacturer recommendations in basal medium (Mesenchymal Stem Cell Basal Medium for Adipose, Umbilical and Bone Marrow-derived MSCs, Cat: PCS-500-030) with supplementation (Mesenchymal Stem Cell Growth Kit for Bone Marrow-derived MSCs, Cat: PCS-500-041) also purchased from ATCC. Cells were maintained at 37°C in an atmosphere of 5% carbon dioxide. Using the human MSC analysis kit (BD Biosciences), cells were found to be conforming to the definition of stemness as outlined by the International Society for Cellular Therapy (43). After two passages using Accutase (STEM CELL Technologies), cells were verified to be mycoplasma free, and stocks were made in 10% DMSO at a concentration of about 1×10^6^ cells per vial. Human umbilical vein endothelial cells (HUVEC), GH354 cells, and murine distal convoluted cells (mDCT) were also purchased from ATCC and grown as per manufacturer’s recommendations in their respective culture media. Antibiotics were not used in any step of the process.

### Synchronized gametocyte culture

*P. falciparum* NF54 parasites were freshly thawed for each experiment from a master stock and cultured in RPMI 1640 (with L-Glutamine, HEPES, and Hypoxanthine) supplemented with 10% O^+^ human serum (Interstate Blood Bank) and 2.5g/L sodium bicarbonate as previously described (44). This was referred to as conventional culture medium, Conv, in this study. Culture medium was replenished daily, and parasites were maintained at 37°C in an environment of 5% oxygen, 5% carbon di-oxide and 90% nitrogen. The first replicative cycle was carried out at 2% hematocrit using O^+^ RBCs (Interstate Blood Bank). As the culture entered the second cycle, it was adjusted to 4% hematocrit and maintained at that level until sufficient parasite volume had been cultured to meet the requirements for each experiment. Once sufficient volume of culture was attained, the culture was synchronized to ring stages using 5% sorbitol solution and this was considered day minus four (D-4). On D-3, trophozoites were diluted to 1% parasitemia at 4% hematocrit. This sorbitol and dilution procedure was repeated on D-2 and D-1. On D0, parasites were at nearly 5% ring parasitemia and 4% hematocrit. At this point, gametocyte induction was performed by adding an equal volume of RPMI 1640 (with L-Glutamine, HEPES, and Hypoxanthine) supplemented with 0.39% fatty acid free BSA, 30μM oleic acid and 30 μM palmitic acid (45); to the existing spent medium in the flask. On D1, the induced culture was spun down and supernatant was completely aspirated before diluting to 1% total parasitemia, 4.5% hematocrit using Conv. On D2, the culture medium was replenished with fresh Conv medium and on D3, co-culture began. No heparin, N-acetyl glucosamine, or antibiotic treatments were performed on these parasites.

### Co-culture of synchronized gametocytes and MSCs

As parasite culture progressed, on D-4, a single vial of MSC cells (1×10^6^) was thawed and used to seed the required number of vented T25 flasks in stem cell medium as described above. Typically, within a week, they attained confluence and were ready to be used for co-culture. Gametocyte culture was divided into as many test groups as required and centrifuged to pellet RBCs and all supernatant was aspirated. The infected RBC pellets were then resuspended in a 1:1 hybrid of Conv medium and stem cell medium (referred to as Hyb in this study) to attain 2% hematocrit. In the initial experiments shown in Fig 1, a part of the infected RBC pellets was resuspended in 100% Conv medium as a negative control. After removing the culture medium in the T25 flasks with MSC cells,10mL of parasite suspension was gently layered onto MSCs for co-culture, or empty T25 flasks for medium controls. The flasks were maintained at 37°C in an environment of 5% oxygen, 5% carbon dioxide and 90% nitrogen. Spent culture medium was aspirated daily and replenished with 7-8 mL (per flask) of freshly prepared Hyb or 100% Conv for Fig 1, or CM for Fig 6 and 7. After 11 days of co-culture (D14), NF54 gametocytes matured to stage V and were ready to infect mosquitos.

### Mosquito infections

*Anopheles stephensi* mosquitos were maintained and feeding experiments were conducted as previously described in the standard membrane-feeding assay (SMFA) protocol (44). At the completion of co-culture, percent stage V gametocytemia was determined by a thin smear, and exflagellation per μL from 25% hematocrit culture was counted on a Nexcelom cellometer for each condition. All test conditions in an experiment were adjusted to the same percentage stage V gametocytemia (0.1 to 0.4 % unless specified otherwise), in 200µL of 50% hematocrit culture and fed to 4–5-day old mosquitos. The percent stage V gametocytemia, exflagellation number and feed gametocytemia in each experiment are shown in Table S1. After 8 days of maintenance, oocysts in the midguts of 20 mosquitoes for each test group were enumerated as previously described (44).

### Co-culture with TNFα treated MSCs

For the experiments where MSCs were pre-treated with TNFα, the cells were incubated with 1ng/mL TNFα (Peprotech) overnight on D2. The next morning supernatant was aspirated, and cells washed twice in Hyb. Co-culture for all conditions proceeded as previously described.

### Co-culture with other mammalian cells

Experiments involving HUVEC, mDCT, and GH 354 cells were carried out in a similar manner to MSC co-culture previously described. The adherent cells were seeded and allowed to reach a confluent monolayer before synchronized gametocytes were layered on them. They were maintained in relevant Hyb.

### Growing gametocytes in substrate coated flasks

For experiments involving substrate coatings, in the morning of D3, empty T25 culture flasks were coated as per manufacturer’s instructions with 9 μg/cm^2^ collagen, 2.5 μg/cm^2^ fibronectin, or 1:100 diluted Matrigel (all from Sigma-Aldrich). Coated flasks were gently washed with Hyb once, then synchronized gametocyte cultures were added to the flasks. The cultures were maintained for 11 days as previously described.

### MSC co-culture of field isolated *P. falciparum*

Two parasites isolated from Cambodian (# 887, and # 806) and Malian patients (KN # 1254, and KN # 1068) were co-cultured in this study with MSCs as minimally culture-adapted *P. falciparum* parasites (46–49). Unlike NF54, these required eighteen days in co-culture to generate mature gametocytes. After this period, the mature gametocytes were used to infect mosquitos by SMFA. For the feeding experiments with the Malian parasites, levels of % stage V gametocytemia among different cultures were not adjusted, as stage V gametocytemia was too low to determine by thin smear.

### Generating conditioned medium

MSCs were grown to confluence in vented T25 flasks. They were then incubated in 10mL Hyb medium for 24 hours and designated as feeder flasks. Supernatant was harvested and filtered through 0.2µm filter daily to generate CM. This procedure was altered to include infected or uninfected RBCs to generate altered conditioned media as described in the associated experiments. CM were harvested from feeder flasks concurrently during experimentation and used fresh in their respective test conditions unless specified. In experiments described in Fig 7, the CM was frozen at -80°C and warmed to 37°C prior to use.

### Statistical analysis

Statistical difference in oocyst numbers between two groups was assessed using a zero inflated negative binomial (ZINB) model described before (50) and calculated by R studio (Ver 1.3.1056). When more than 3 groups were compared, Bonferroni adjustments were performed. To compare average oocyst numbers from ten independent experiments with or without MSC CC, Wilcoxon matched pairs signed rank test was used. *P*< 0.05 was considered significant. Data were visualized using GraphPad Prism (Ver 9.3.1)

### Ethics statement

Field parasites were isolated from patients after obtaining written, informed consent in prior studies (46–49). The original epidemiology studies were approved by the Cambodian National Ethics Committee for Health Research, the Ethics Committee of the Faculty of Medicine, Pharmacy and Odontostomatology at the University of Bamako and the National Institute of Allergy and Infectious Diseases Institutional Review Board (ClincalTrials.gov identifier: NCT00341003, and NCT00669084).

## Acknowledgements

This work was supported by the intramural program of the NIAD/NIH. Our thanks to Drs. Juliana Sa, and Thomas Wellems for sharing field isolated parasites.

We also thank Dr. Francesco Silvestrini for graciously sharing the initial protocol for growing gametocytes in the presence of MSCs which was extensively modified to reach the protocol presented in this study.

## Notes

### Competing Interest Statement

The authors have declared no competing interest.

