## Supplemental Fig 1, 2 and table 1 for "Mesenchymal stem cells of the bone marrow raise infectivity of *Plasmodium falciparum* gametocytes"

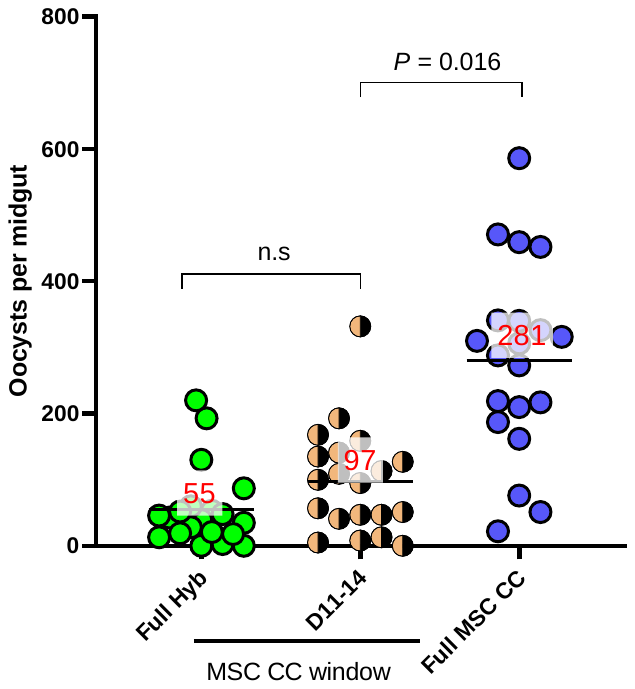

**Fig. S1.** **Co-culture for the last four days of gametocyte development did not significantly raise oocyst yields relative to Hyb control.** *P. falciparum* NF54 gametocytes were co-cultured with MSCs for D11 to D14 and full duration (D3-14, Full MSC CC) or full duration Hyb medium. These gametocytes were used to infect mosquitos and resulting oocysts per midgut are presented with the arithmetic means (bars with red text). P value was calculated by ZINB. (n.s) no significant difference. The Full Hyb and Full MSC CC data in this figure are the same as shown in Fig 4.

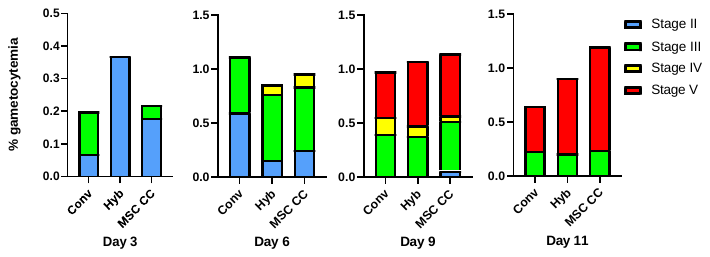

**Fig. S2.** **Gametocytes undergo similar developmental stages during co-culture.** *P. falciparum* NF54 gametocytes were co-cultured with human mesenchymal stem cells, (MSC CC), in conventional gametocyte culture medium (Conv), or in hybrid culture medium (Hyb). Gametocytemia was measured by Giemsa staining thin smears of culture. The percent of gametocyte stages measured on days 3, 6, 9, and 11 with bars representing the median values from three cultures per group in a single experiment.

**Table S1:Culture parameters on the day of feeding experiment, and final feed gametocytemia:**

| Figure # | Condition | % Stage V gametocytemia | Exflagellation per μL | % Stage V gametocytemia fed to mosquitos |
| --- | --- | --- | --- | --- |
| 1B | Conv | 1.94 | 1285 | 0.40 |
|  | Hyb | 2.98 | 620 |  |
|  | MSC CC(+TNFα) | 1.82 | 1000 |  |
| 1D | Conv | 1.16 | 545 | 0.11 |
|  | Hyb | 0.74 | 50 |  |
|  | MSC CC(+TNFα ) | 0.68 | 580 |  |
| 2A | Hyb | 2.28 | 200 | 0.11 |
|  | MSC CC(+TNFα) | 1.64 | 1280 |  |
|  | MSC CC | 1.76 | 690 |  |
| 3A | HUVEC Hyb | 0.16 | n.d. | 0.20 |
|  | HUVEC CC | 0.76 | 70 |  |
|  | MSC Hyb | 1.04 | 40 |  |
|  | MSC CC | 0.91 | 240 |  |
| 3B | GH354 Hyb | 0.50 | 780 | 0.20 |
|  | GH354 CC | 0.80 | n.d. |  |
|  | MSC Hyb | 1.32 | 710 |  |
|  | MSC CC | 0.84 | 1550 |  |
| 3C | mDCT Hyb | 1.05 | 10 | 0.20 |
|  | mDCT CC | 0.71 | n.d. |  |
|  | MSC Hyb | 1.04 | 40 |  |
|  | MSC CC | 0.91 | 240 |  |
| 3D | Fibronectin | 1.51 | 380 | 0.11 |
|  | Matrigel | 0.99 | 270 |  |
|  | Collagen | 1.17 | 240 |  |
|  | Hyb | 1.21 | 290 |  |
|  | CC | 0.91 | 370 |  |
| 4A | Full Hyb | 1.67 | 150 | 0.10 |
|  | [D3-6] | 0.82 | 260 |  |
|  | [D3-10] | 0.76 | 270 |  |
|  | MSC CC | 1.06 | 150 |  |
| 5A | Hyb | 0.35 | n.d. | 0.15 |
|  | MSC CC | 0.44 | 70 |  |
| 5B | Hyb | 0.05 | n.d. |  |
|  | MSC CC | 0.04 | n.d. |  |
| 5C | Hyb | n.d. | n.d. | 120 µL of packed RBC from each culture were fed without adjustment of Stage V gametocytemia |
|  | MSC CC | n.d. | n.d. |  |
| 5D | Hyb | n.d. | n.d. |  |
|  | MSC CC | n.d. | n.d. |  |
| 6A | Hyb | 0.70 | 70 | 0.40 |
|  | CM | 1.81 | 160 |  |
|  | MSC CC | 0.31 | 330 |  |
| 6B | Hyb | 1.59 | 260 | 0.16 |
|  | CM (MSC^+^ iRBC^-^) | 1.38 | 270 |  |
|  | CM (MSC^-^ iRBC^+^) | 0.88 | 180 |  |
|  | CM (MSC^+^,iRBC^+^) | 0.73 | 150 |  |
|  | MSC CC | 1.54 | 970 |  |
| 6C | Hyb | 0.91 | 310 | 0.1 |
|  | CM (MSC^+^ uRBC^-^) | 0.31 | 340 |  |
|  | CM (MSC^-^ uRBC^+^) | 0.35 | 140 |  |
|  | CM (MSC^+^,uRBC^+^) | 0.3 | 480 |  |
|  | MSC CC | 0.56 | 800 |  |
| 7A | Hyb | 0.24 | 90 | 0.15 |
|  | CM | 0.32 | 50 |  |
|  | CM (Frozen) | 0.26 | 50 |  |
|  | MSC CC | 0.25 | 320 |  |
| 7B | Hyb | 0.44 | 300 | 0.15 |
|  | CM | 0.62 | 440 |  |
|  | CM (Frozen) | 0.53 | 520 |  |
|  | MSC CC | 0.32 | 220 |  |
| S1 | Hyb | 1.67 | 150 | 0.10 |
|  | [D8-11] | 0.93 | 970 |  |
|  | MSC CC | 1.06 | 150 |  |

n.d. : not detectable (<0.001% gametocytemia or <2.5 exflagellation center per µL)
